# Single Cell Molecular Alterations Reveal Pathogenesis and Targets of Concussive Brain Injury

**DOI:** 10.1101/250381

**Authors:** Douglas Arneson, Yumei Zhuang, Hyae Ran Byun, In Sook Ahn, Zhe Ying, Guanglin Zhang, Fernando Gomez-Pinilla, Xia Yang

**Affiliations:** Department of Integrative Biology and Physiology, University of California, Los Angeles, Los Angeles CA 90095, USA.; Bioinformatics Interdepartmental Program, University of California, Los Angeles, Los Angeles CA 90095, USA.; Department of Neurosurgery, University of California, Los Angeles, Los Angeles CA 90095, USA.; Institute for Quantitative and Computational Biosciences, University of California, Los Angeles, Los Angeles CA 90095, USA.; Molecular Biology Institute, University of California, Los Angeles, Los Angeles CA 90095, USA.

## Abstract

The complex neuropathology of traumatic brain injury (TBI) is difficult to dissect in the hippocampus considering the convoluted hippocampal cytoarchitecture. As a major casualty of TBI, hippocampal dysfunction results in cognitive decline that may escalate to other neurological disorders, and the molecular basis is hidden in the genomic programs of individual hippocampal cells. Using the unbiased single cell sequencing method Drop-seq, we uncovered the hippocampal cell types most sensitive to concussive mild TBI (mTBI) as well as the vulnerable genes, pathways and cell-cell interactions predictive of disease pathogenesis in a cell-type specific manner, revealing hidden pathogenic mechanisms and potential targets. Targeting *Ttr,* encoding the thyroid hormone T4 transporter transthyretin, mitigated the genomic and behavioral abnormalities associated with mTBI. Single cell genomics provides unique evidence about altered circuits and pathogenic pathways, and pinpoints new targets amenable to therapeutics in mTBI and related disorders.

---

TBI is common in domestic, sports, and military environments and often leads to long-term neurological and psychiatric disorders^1^. The hippocampus is a member of the limbic system and plays a major role in learning and memory storage. As a major aspect of the TBI pathology^2^, hippocampal dysfunction leads to memory loss and related cognitive impairment. The hippocampal formation encompasses four Cornu Ammonis (CA) subfields largely composed of pyramidal cells, and their connections with dentate gyrus cells. The CA - dentate gyrus circuitry has served as a model to study synaptic plasticity underlying learning and memory. Glial cells are vital to the hippocampal cytoarchitecture, however, their interactions with neuronal cells are poorly defined. The heterogeneous properties of the hippocampal cytoarchitecture have limited the understanding of the mechanisms involved in the TBI pathology. Mild TBI (mTBI) is particularly difficult to diagnose given its broad pathology, such that there are no accepted biomarkers for mTBI^3^. This limitation becomes an even more pressing issue given the accumulating clinical evidence that mTBI poses a significant risk for neurological and psychiatric disorders within the spectrum of the hippocampus such as Alzheimer’s disease (AD), chronic traumatic encephalopathy (CTE), epilepsy, and dementia^4^. Accordingly, there is an urgent need to identify functional landmarks with predictive power within the hippocampus to address current demands in clinical neuroscience.

Given that gene regulatory programs determine cellular functions, scrutiny of large-scale genomic changes can reveal clues to the molecular determinants of mTBI pathogenesis including cellular dysfunction, injury recovery, treatment response, and disease predisposition. However, existing genomic profiling studies of mTBI are based on heterogeneous mixtures of cell conglomerates^5-9^ which mask crucial signals from the most vulnerable cell types. Here, we report the results of a high throughput parallel single cell sequencing study, using Drop-seq, to capture mTBI-induced alterations in gene regulation in thousands of individual hippocampal cells in an unbiased manner. We focused on concussive injury, the most common form of mTBI, using a mild fluid percussion injury (FPI) mouse model which induces identifiable hippocampal-dependent behavioral deficits despite minimal cell death^10^. We examined the hippocampus at 24h post-mTBI, as this is a pivotal timeframe for pathogenesis and is generally used for diagnostic and prognostic biomarker discovery^11^.

To our knowledge, this is the first parallel single cell sequencing study to investigate the mTBI pathogenesis in thousands of individual brain cells, offering a cell atlas of the hippocampus under both physiological and pathological conditions. In doing so, we provide novel evidence about the cellular and molecular remodeling in the hippocampus at the acute phase of TBI, and help answer critical longstanding questions. Which cell types are the most vulnerable to mTBI at the acute phase? Within each cell type, which genes have altered transcriptional activities that are induced by mTBI? Which molecular pathways are perturbed by mTBI in each cell type and how do they relate to mTBI pathology and pathogenesis of secondary brain disorders such as AD and PTSD? Which cell-cell communications are disrupted in mTBI? Through answering these questions, the identified sensitive cell types and associated gene markers can serve as signatures of mTBI pathology that inform on the stage, functional alterations, and potential clinical outcomes. Since the cell is the elementary unit of biological structure and function, we reveal fundamental information that can lead to a better understanding of the mechanistic driving forces for mTBI pathogenesis and identify potential novel therapeutic targets in an unbiased manner. As a proof of concept, we used the data-driven single cell information to prioritize *Ttr*, encoding transthyretin, as a plausible target and show for the first time that modulating *Ttr* improves behavioral phenotypes and reverses the molecular changes observed in mTBI.

## RESULTS

### Unbiased identification of cell identities in hippocampus

Using Drop-seq^12^, we sequenced 2,818 and 3,414 hippocampal cells from mTBI and Sham animals, respectively. A single-cell digital gene expression matrix was generated using Drop-seq Tools^12^ and subsequently projected onto two dimensions using t-distributed stochastic neighbor embedding (t-SNE)^13^ to define cell clusters (**Methods**). We detected 15 clusters each containing cells sharing similar gene expression patterns (**Fig. 1a**). The cell clusters were not due to technical or batch effects (**Supplementary Fig. 1**).

**Figure 1.**
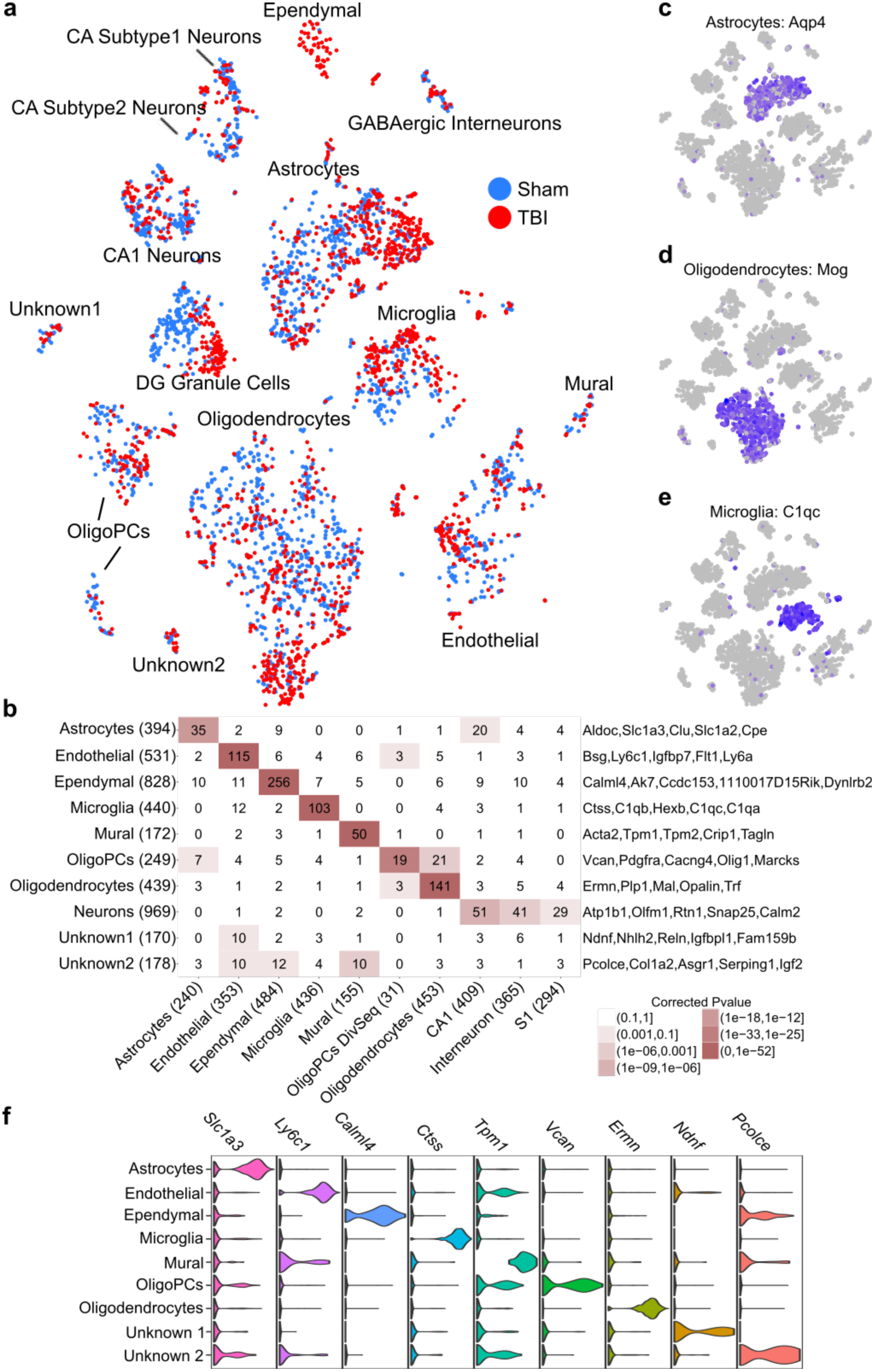
Determination of major hippocampal cell types and cell type-specific gene markers using the unbiased Drop-seq approach. (**a**) t-SNE plot showing cell clusters. Each colored dot is a cell, with blue cells originating from Sham animals and red cells originating from mTBI animals. (**b**) Overlap between Drop-seq defined marker genes of major cell clusters (rows) with known cell type markers (columns) derived from a previous Fluidigm-based single cell study^14^. Signature marker numbers are indicated in the parenthesis. Statistical significance of overlap is indicated by color (the darker the more significant) and the numbers of overlapping genes between our Drop-seq defined markers, and previously known markers are shown in the cells. Top cell marker genes determined by our Drop-seq data are listed on the right of the plot. (**c-e**) Cluster-specific expression of known cell markers: Astrocytes - *Aqp4*, Oligodendrocytes – *Mog*, and Microglia – *C1qc*. This analysis confirms that each cluster captures a particular cell type. (**f**) Normalized expression values of top cell type-specific marker genes are plotted as violin plots with cell types as rows and genes as columns.

To resolve the cell-type identities, we obtained cluster-specific gene signatures (**Supplementary Table 1**) and compared them to known signatures of hippocampal cell types derived from Fluidigm-based single cell studies^14, 15^ (**Methods**). We recovered all known major cell types including neurons, oligodendrocytes, microglia, mural cells, endothelial cells, astrocytes, and ependymal cells (**Fig. 1b**). Previously known cell markers, such as *Aqp4* for astrocytes, *Mog* for oligodendrocytes, and *C1qc* for microglia, all showed distinct cluster-specific expression patterns, confirming the reliability of our data-driven approach in distinguishing cell types (**Fig. 1c-e**; additional known marker examples in **Supplementary Fig. 2**). Beyond retrieving known cell markers, our Drop-seq paradigm also enabled us to identify novel marker genes for each cell type, such as *Calml4* for ependymal, *Vcan* for oligodendrocytes, and *Ly6a* for endothelial cells (**Fig. 1f**; **Supplementary Table 1**). We confirmed the cellular identity and localization of many novel marker genes using the *in situ* hybridization (ISH) images from the Allen Brain Atlas (**Fig. 2**; **Supplementary Fig. 3-9**).

**Figure 2.**
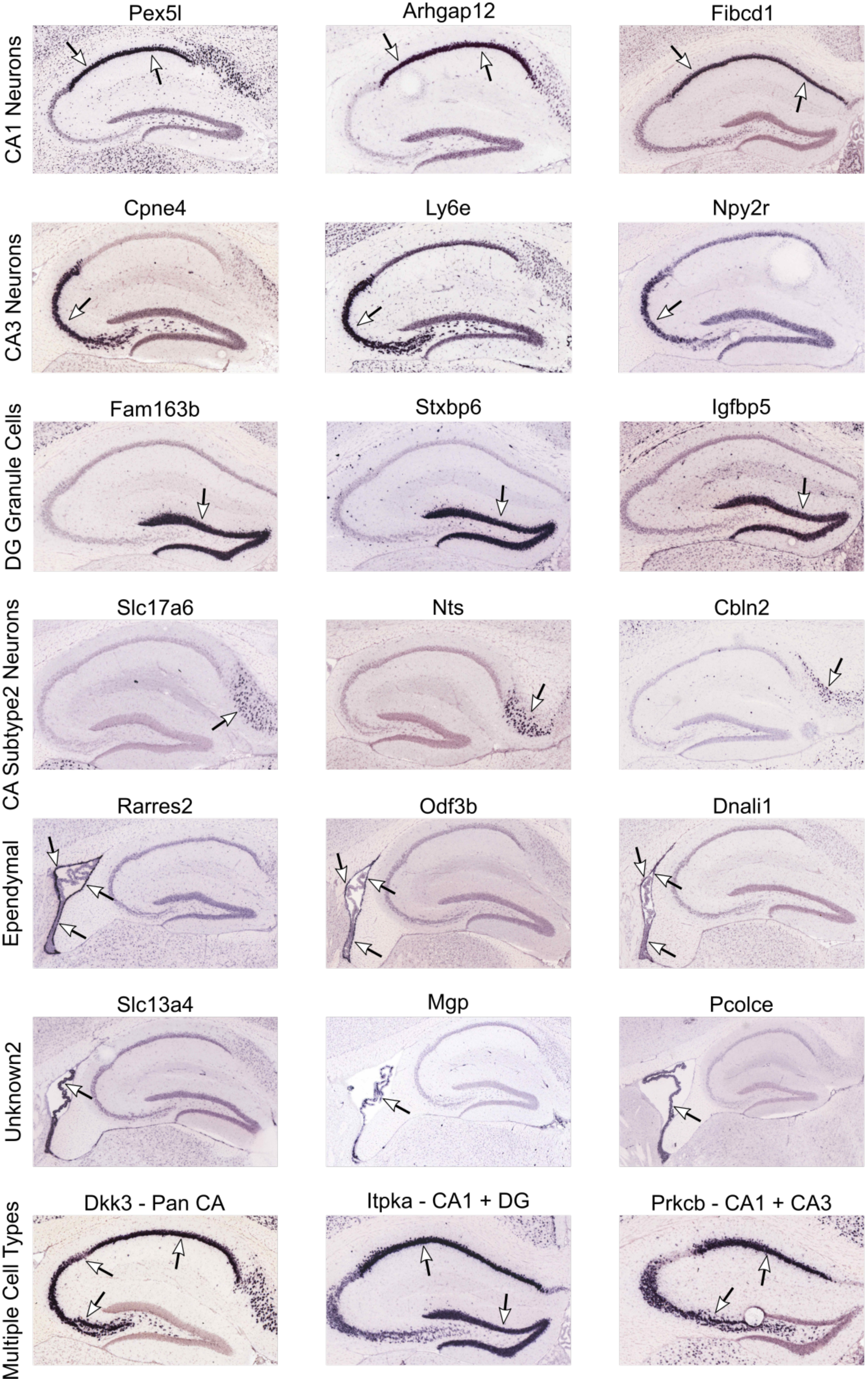
Cross-validation of novel marker genes for specific neuronal subpopulations. To validate the specificity of novel marker genes for neuronal populations and to help resolve the identity of previously unknown cell clusters, we examined the expression patterns of our cell markers in the ISH images from the Allen Brain Atlas^56^. Here, we showcase three select novel genes from four cell types: CA1 neurons, CA3 neurons, DG granule cells, and ependymal cells. Additionally, we showcase marker genes expressed across multiple cell types, genes which clearly resolve the Unknown2 cluster to cells inside the choroid plexus, and genes which help resolve CA Subtype2 Neurons to the Subicular Complex.

We identified two novel cell clusters whose gene expression signatures did not significantly overlap with any of the known cell types that we were expecting in the hippocampus, and were hence named “Unknown1” (totaling 113 cells and representing 1.8% of all cells tested) and “Unknown2” (totaling 74 cells and representing 0.9% of all cells examined; **Fig. 1a**). Unknown1 has markers indicative of cell growth and migration such as *Ndnf, Nhlh2*, *Reln*, and *Igfbpl1*, as well as endothelial markers (**Fig. 1b**), suggesting that these clusters may represent migrating endothelial cells. This is consistent with the role of endothelial cells as main components of the blood brain barrier which is disrupted after mTBI^16^, with proliferation and migration of endothelial cells being an intrinsic aspect of new vessel formation^17^. Unknown2 expresses unique markers indicative of cell differentiation such as *Pcolce*, *Col1a2*, *Asgr1*, *Serping1*, and *Igf2*, along with markers of endothelial, mural, and ependymal cells (**Fig. 1b**), suggesting that these may be progenitor cells differentiating into multiple lineages. To further resolve the identities of these unknown clusters, we examined the expression patterns of their signature genes identified by our single cell sequencing in the ISH images in the Allen Brain Atlas. Using this method, the marker genes of the Unknown2 cluster colocalized to a population of cells in the choroid plexus distinct from the ependymal cells (**Fig 2**; **Supplementary Fig. 8**). This localization to an area known to house stem cells and progenitor cells taken in conjunction with the functions of the marker genes strengthens the claim that this cell population may represent progenitor cells. These unknown cell clusters illustrate cell types that have been left undetected by classical morphological categorization, and may represent potentially novel functional entities within the hippocampal formation. Additionally, we were able to categorize oligodendrocyte progenitor cells (**Fig. 1a**) which were undetected in previous lower-throughput single cell studies^14, 15^.

To further characterize neuronal diversity within the hippocampus, we detected nine neuronal subclusters using the BackSPIN biclustering method^14^ (**Fig. 3a**). Annotation with known neuronal markers helped resolve GABAergic interneurons, dentate gyrus (DG) granule cells, and 4 subtypes of CA pyramidal neurons (**Fig. 3b**). However, two clusters (“Neuronal Subtype1”, “Neuronal Subtype 2”) remained unannotated, representing potential novel neuronal subtypes or states. Based on the functions of their markers (**Fig. 3b**) it is possible that these clusters may contain neurons with the potential to differentiate or self-renew. The unique ability of Drop-seq to catalog cells based on unbiased genomic parameters was also instrumental in unveiling novel neuronal markers^14, 15^ including *Ptn* in CA1 neurons, *Ly6e* in CA3 neurons, and *Sema5a* in DG granule cells (**Fig. 3b**; full marker list in **Supplementary Table 1**). We confirmed the subtype-specific expression patterns in specific hippocampal subregions and cell types (**Fig. 3c-e**) and further verified the expression specificity of several novel markers using ISH data from the Allen Brain Atlas (**Fig. 2, Supplementary Fig. 3-9**). The Allen Brain Atlas ISH data also helped resolve the identity of the CA Subtype2 cells to be from the Subicular Complex, which is located proximal to the CA1 region (**Fig. 2**) and mediates the main output of signals from the hippocampus. Like the CA subregions, the Subicular Complex is also comprised of pyramidal neurons, which explains the expression of CA neuronal signature genes in addition to the genes specific to this cell cluster. It has been reported that CTE and associated dementia involves damage to the subiculum^4^, which is in general agreement to the proposed involvement of subiculum in the pathology of Alzheimer’s Disease^18^.

**Figure 3.**
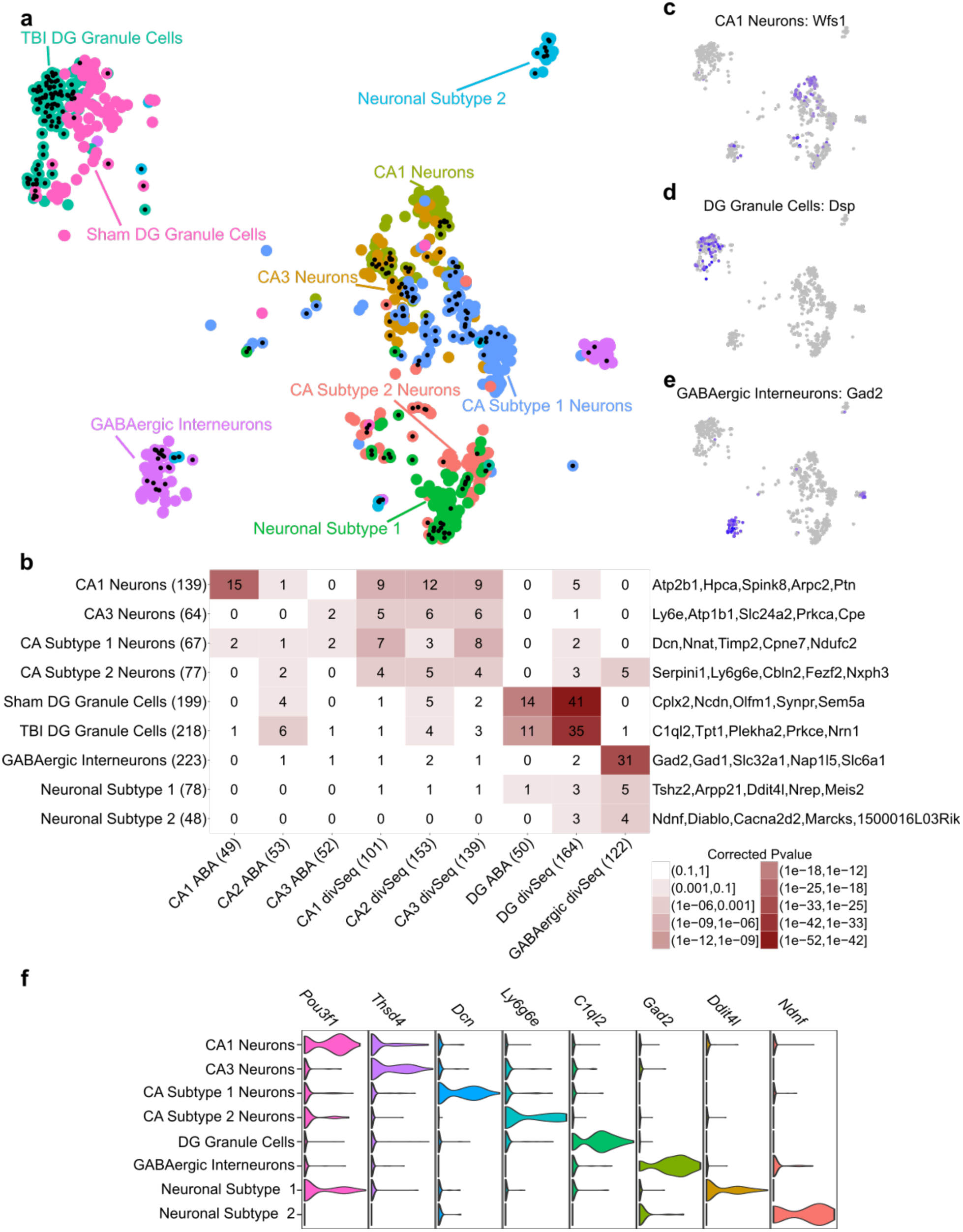
Determination of neuronal cell subtypes and cell type-specific gene markers using the unbiased Drop-seq approach. (**a**) t-SNE plot of neuronal subtypes determined by backspin biclustering. Each color indicates a different cell type cluster identified, and cells with a black dot at their center from TBI samples. (**b**) Overlap of Drop-seq defined marker genes of the neuronal subtypes (rows) with those of the previously defined hippocampal neuronal cell types (columns). Known markers were derived from Alan Brain Atlas (ABA)^56^ and Habib et al. using Div-Seq^15^. Signature marker numbers are indicated in the parenthesis. Statistical significance of overlap is indicated by color (the darker the more significant), and the numbers of overlapping genes between our Drop-seq defined markers and previously known markers are shown in the cells. Top cell marker genes determined by our Drop-seq data are listed on the right of the plot. **(c-e**) Cluster-specific expression of known cell markers: CA1 neurons – *Wfs1*, DG granule cells – *Dsp*, and GABAergic interneurons – *Gad2*. (**f**) Normalized expression values of top neuronal subtype-specific marker genes are plotted as violin plots with cell types as rows and genes as columns.

These results indicate that our transcriptome-driven, unbiased Drop-seq approach has the unique ability to uncover new cell types, states, and markers based on genomic features that determine function, an undertaking not possible with traditional morphology-based approaches or bulk tissue sequencing. These novel findings provide valuable resources for future studies of the hippocampal circuitry under homeostatic and/or pathological conditions.

### Identification of most vulnerable cell types to mTBI based on transcriptomic profile

To retrieve hippocampal cell types vulnerable to mTBI, we first compared the cell population proportions between mTBI and Sham animals. We found that ependymal cells are more abundant in mTBI compared to Sham animals (89% of ependymal cells from mTBI samples vs 11% from Sham samples, **Fig. 1a**). This large shift in relative abundance of ependymal cells at 24 h post-surgery implicates them as key acute responders to mTBI. This is consistent with the reported role of ependymal cells in acute post-injury processes such as circuit repair and scar formation^19, 20^. In addition to the visible increase in the relative abundance of ependymal cells in mTBI, DG granule cells are clearly separated into two clusters, one almost exclusively (94%) from the Sham animals and another primarily (86%) from the mTBI samples (**Fig. 1a, Fig. 3a**). Post-traumatic epilepsy is a major concern in TBI and is attributable to DG dysfunction^21^. Other hippocampal cell types that demonstrate weaker yet visible transcriptomic shifts between mTBI and Sham include oligodendrocytes, astrocytes, microglia, and multiple neuronal subpopulations such as CA1 neurons and CA subtype2 neurons clusters (**Fig. 1a**, **Fig. 3a**).

The identification of specific cell types particularly sensitive to mTBI, like the ependymal cells, astrocytes, and neuronal clusters such as DG granule cells and CA pyramidal cells, opens the possibility for investigating the specific roles of each cell type in mTBI pathology and for designing targeted treatments. For example, the genomic markers of the DG granule cells highly sensitive to mTBI can be used for the design of treatments that target specific DG subpopulations responsible for post-traumatic epilepsy to avoid the side effects of classical epilepsy treatments targeting broad populations of cells.

### mTBI disrupts cell-cell interaction within the hippocampal circuitry

Emerging evidence in the neuroimaging field suggests that changes in the interaction patterns among cells in circuits can contribute to reduced cognitive capacity in TBI^22^. We used the co-regulation patterns between genes of different cell types to infer interactions among cells, as gene co-expression can infer functional connectivity^23, 24^. Specifically, we focused on marker genes encoding secreted peptides from each cell type (source cells), which have the potential to interact with genes in other cell types (target cells) (**Methods**). This gene co-expression analysis showed extensive reorganization in the pattern of interaction among cells in response to mTBI. For example, the interaction from astrocytes and ependymal cells to neurons and from microglia to oligodendrocytes was enhanced in mTBI (**Fig. 4**). We also found in mTBI, decreased interaction between microglia and neurons, and decreased interaction from oligodendrocytes to neurons. These shifts in the pattern of interactions may reflect a unique and novel property of hippocampal cells to reorganize the working flow in response to mTBI challenge. The changes in the interaction were also reflected in changes in the association pattern among single genes. For instance, the correlation of *Bdnf* from neurons with cell metabolism genes in microglia was lost in mTBI. Notably, only in mTBI, the known AD risk gene *Apoe* in astrocyte and ependymal cells (source cells) showed strong correlations with mitochondrial metabolism genes in neurons (target cells). These results suggest the potential of mTBI to alter the interactive patterns at the level of circuits and genes in the hippocampus, with cell metabolism involved in the interactions.

**Figure 4.**
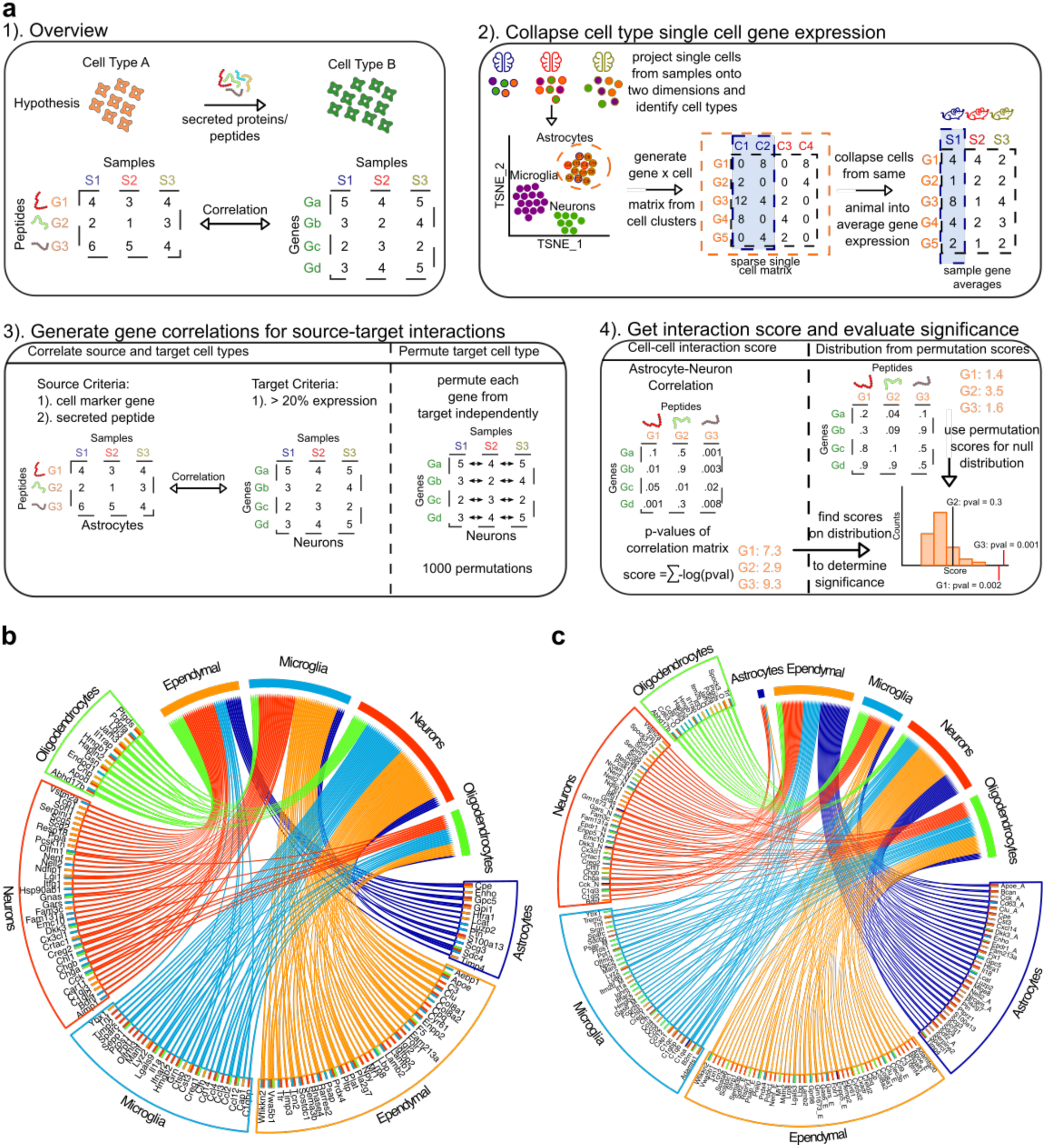
TBI alters cell-cell interations, and promotes molecular reorganization in hippocampus. (**a**). Methodology used for cell-cell interaction analysis. Secreted proteins or peptides from a source cell can communicate with genes in a target cell, which can be captured by strong correlations between the secreted proteins in the source cells and genes in the target cells. For each cell cluster, the expression of each gene was summarized to individual TBI or Sham animal level, and a correlation matrix between genes of different cell types was constructed. The −log10 p-values of these correlations for each secreted peptide are summed to obtain an interaction score of that peptide with a particular target cell type. The cell type gene expression matrix is then permuted to generate the null distribution of interaction scores to calculate the significance of an observed interaction score. Significant interactions are plotted in the circos plots for the Sham group (**b**) and the TBI group (**c**) separately. The bottom half of each circos plot shows source cell types with secreted peptides (names listed) and the top half are the target cell types. Colored lines in the center indicate significant connections between the peptides with different cell types. Comparison between (**b**) and (**c**) shows reorganization in the interactions among different cell types within the hippocampal formation after TBI and the associated genes potentially mediating the interactions. As the transcriptome of individual cells can instruct cell-cell communication to process high order information, these results suggest potential changes in neural circuit organization after an episode of TBI.

### Identification of genes and pathways vulnerable to mTBI in individual cell types

To determine specific genes and pathways that may confer mTBI pathogenesis in each cell type and to further refine the vulnerable cell types, we identified differentially expressed genes (DEGs) between Sham and mTBI within each cell cluster (**Table 1**; all DEGs in **Supplementary Table 2**) at false discovery rate (FDR) <0.05. Astrocytes, oligodendrocytes, and neurons had the largest numbers of DEGs between mTBI and Sham samples (**Table 1**). Annotation of the DEGs with curated biological pathways from KEGG^25^, Reactome^26^, BIOCARTA, and Gene Ontology^27^ databases identified key pathways that could explain fundamental aspects of the mTBI pathology and the main cell types involved (**Table 1**; full pathway list in **Supplementary Table 3**). For example, the DEGs were enriched for diverse pathways encompassing energy and metabolism (astrocytes, neurons), inflammation and immune response (microglia, oligodendrocyte PCs), myelination (oligodendrocytes), amyloids (endothelial and ependymal cells), neurogenesis and synaptic signaling (neurons), cell migration (GABAergic interneurons), glutamate transport (CA1 pyramidal cells), and dendrite morphogenesis (“Unknown 1” cluster). Many of the pathways identified agree with the known roles of the specific cell types in mTBI. For example, the over-representation of inflammation and immune pathways in microglia is consistent with the role of microglia in inflammatory processes and the efficacy of anti-inflammatory treatments targeting microglia at one-day post-mTBI^28^. Nevertheless, the identified cell type specific genes and pathways vulnerable to mTBI also offered novel insights into the functions of individual cell types in mTBI pathogenesis, as detailed below.

**Table 1.**
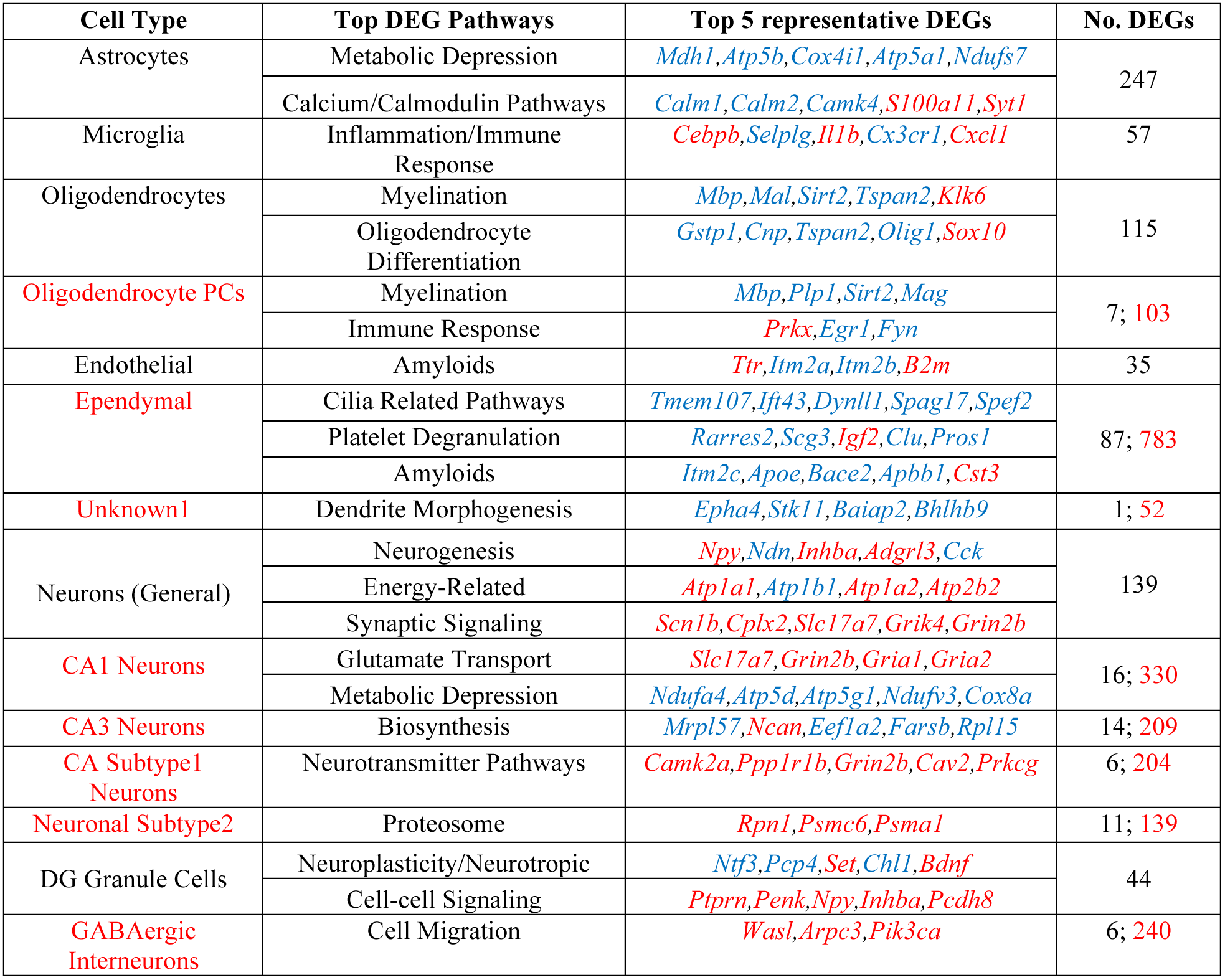
Top enriched pathways among DEGs of major cell types (FDR<5%) and representative DEGs in the select pathways. DEGs in blue and red show decreased and increased expression in TBI, respectively. Cell types shown in red are rare cell types with fewer cells analyzed, and hence did not have sufficient numbers of DEGs at FDR<5% (number of DEGs shown in black) between Sham and TBI samples to reveal significant pathways. Instead, DEGs reaching a threshold of p < 0.01 (number of DEGs shown in red) were used to derive suggestive pathways for these rare cell types.

TBI is followed by a stage of metabolic dysfunction^29^ which reduces the capacity of the brain for activity-dependent plasticity ^30^; however, the molecular and cellular underpinnings remain poorly understood. Consistent with the metabolic crisis typical of mTBI, we found down-regulation of mitochondrial metabolic genes in astrocytes and CA1 pyramidal cells. Astrocytes supply energy substrates to neurons and are essential for neuronal function. Interestingly, our results indicate that CA1 pyramidal cells also experience metabolic suppression at this early stage of mTBI pathogenesis (24 h). In addition, DEGs in CA1 pyramidal cells informed on increased expression of various glutamate transporters, which could explain the altered capacity to sustain long-term potentiation after TBI^31^.

The risks posed by mTBI on development of other neurological disorders is a pressing issue in clinical neuroscience; however, the molecular and cellular bases for this occurrence remain undetermined. DG granule cells, which function to interact with CA pyramidal cells, showed alterations in genes involved in cell-cell signaling (*Npy, Penk, Ptprn, Ihnba*) as well as neuroplasticity genes such as *Bdnf* and *Ntf3*. These pathways may underlie the mTBI sensitivity of DG granule cells and post-traumatic epilepsy occurring after mTBI. Pathways informed by DEGs from the endothelial and ependymal cells implicate the importance of these cell types in the potential for amyloid buildup, as indicated by the upregulation of the known pro-amyloid deposition gene *B2m* and down-regulation of inhibitors of beta-amyloid aggregation and fibril deposition (*Apoe*, *Itm2a, Itm2b,* and *Itm2c*) in these cell types. As dysregulated metabolism and amyloid deposition are key features in AD, CTE, and PD^32^, our study provides detailed information on the specific cell types such as astrocytes, CA1 pyramidal cells, endothelial and ependymal cells that could be the starting loci for the wave of post-mTBI amyloid buildup in chronic mTBI.

### mTBI alters gene expression programs in a cell-type specific manner

We found that many of the DEGs were significantly altered by mTBI in only one cell type (**Fig. 5a-b**) and showed clear cell type-specific shifts in expression patterns (**Fig. 5c-e**), supporting the capacity of mTBI to modulate gene expression in a cell-type specific manner. Importantly, >50% of the DEGs identified at the single cell level were masked in bulk tissue analysis (**Fig. 5f**; bulk tissue-level DEGs in **Supplementary Table 2**). Notably, the unique cell-level DEGs were primarily from cell types of low abundance such as neurons, and the common DEGs between single-cell and tissue-level analyses were mainly from abundant cell types such as astrocytes and oligodendrocytes. These findings highlight the value of extracting genomic information in individual cell types that otherwise would be masked in bulk tissue studies.

**Figure 5.**
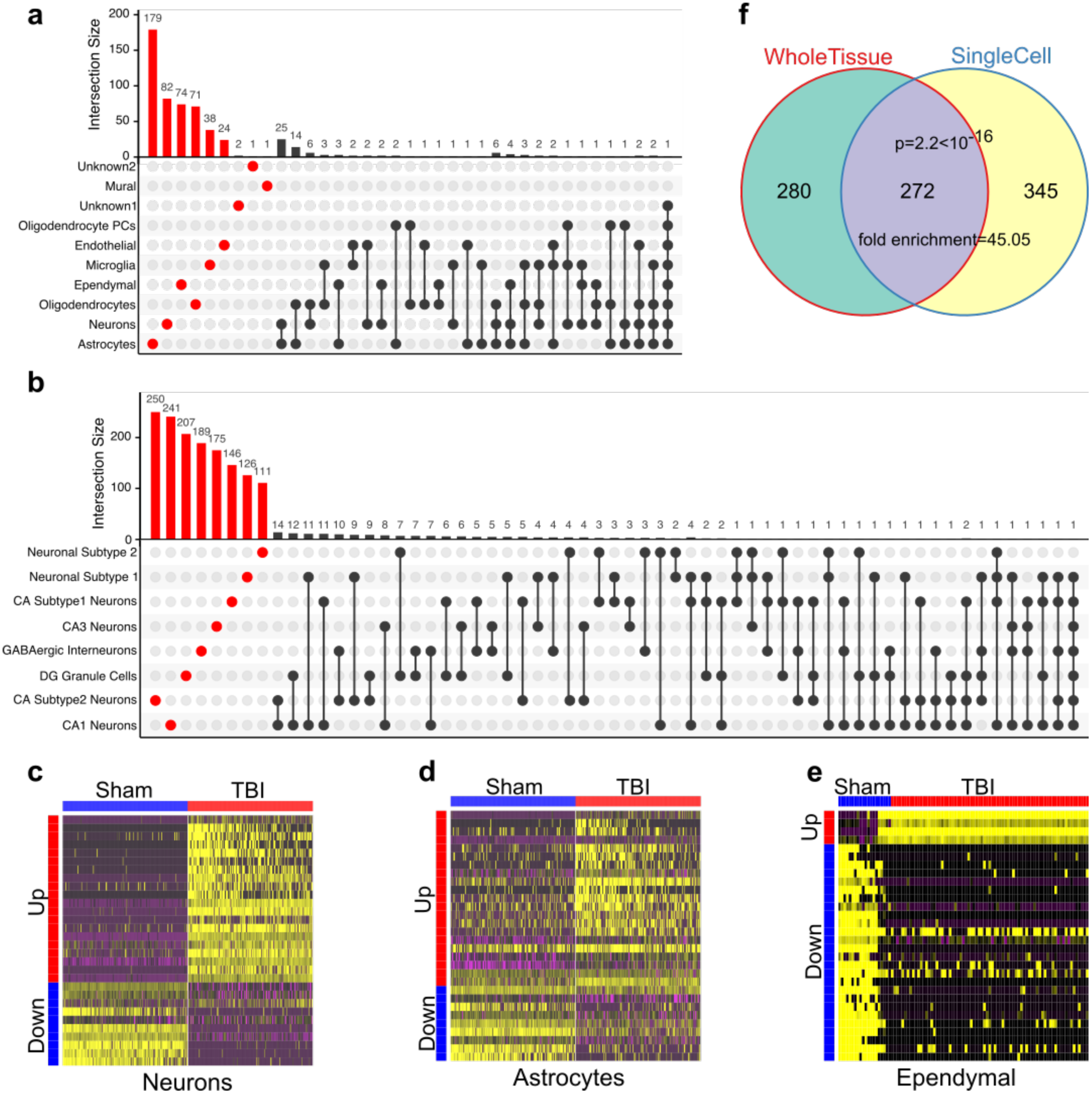
Differentially expressed genes (DEGs) induced by TBI in individual hippocampal cell types identified using Drop-seq. (**a-b**) DEGs unique to a cell type are indicated in red and those shared between >=2 cell types are indicated by black dots. The histogram above each plot indicates the DEG counts for each category. **(a**) The majority of the DEGs are cell-type specific. (**b**) The majority of the DEGs in neuron subtypes are subtype-specific. (**c-e**) Heatmaps of DEGs in select cell types demonstrate clear differential expression patterns between Sham and TBI cells. (**f**) Many cell-type specific DEGs cannot be captured in the bulk tissue analysis, supporting the uniqueness of using single cell genomic analysis.

Cell-type specific DEGs may serve as selective biomarkers or therapeutic targets that can trace or normalize specific abnormalities of mTBI pathology (**Figure 6a**). For instance, *Id2*, a gene previously described to be upregulated by seizures in the DG^33^, is specifically upregulated by mTBI in DG granule cells and thus could serve as a potential target to temper post-traumatic epilepsy after TBI. We also found that *P2ry12*, a gene previously reported as a marker for brain resident microglia^34^, is specifically downregulated in microglia after mTBI, suggesting the potential utility of *P2ry12* to be used as a marker of the early inflammatory response to TBI. Several of the cell-type specific DEGs are related to amyloid deposition and AD, including *Apoe* - an ependymal-specific DEG and known for its effects on AD and TBI^35^, and *Itm2a* – an endothelial specific DEG and an inhibitor of amyloid-beta production and deposition^36^. These results indicate that the putative action of TBI in amyloid deposition involves different genes across different cell types. Interestingly, *Trf,* encoding transferrin for iron transport was upregulated in oligodendrocytes, suggesting a possible involvement of *Trf* on the described association between iron deposition and cognitive deficits in mTBI^37, 38^.

**Figure 6.**
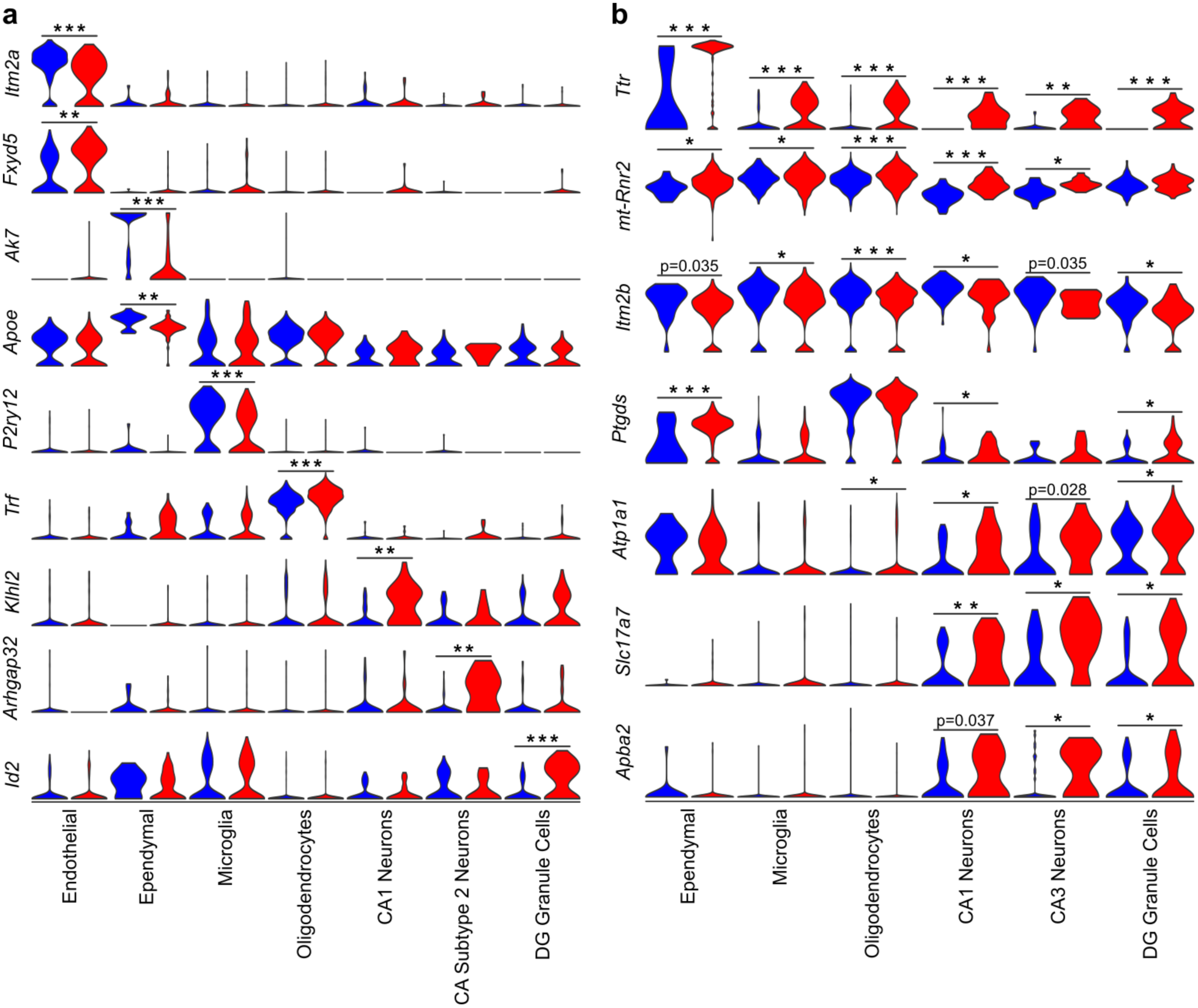
Top cell-type specific DEGs (**a**) and pan-hippocampal DEGs (**b**). The normalized expression of cell-type specific and pan-hippocampal DEGs between Sham and TBI samples is displayed as violin plots. Single cells from Sham samples are indicated by the blue plots and single cells from TBI samples are indicated by the red plots. *p<0.02, **p<1x10^-4^, ***p<1x10^-6^.

We also found several cell-type specific mTBI target genes that have not been implicated in mTBI previously. For instance, a CA1 pyramidal cell-specific DEG *Klhl2*, which encodes an actin binding protein, was recently implicated in human neuroticism^39, 40^. Increased neuronal *Klhl2* expression post-mTBI (**Fig. 6a**) was confirmed at the protein level in the CA1 hippocampal subregion using immunohistochemistry (**Supplementary Fig. 10)**. *Khlh2* in CA1 pyramidal cells may serve as a novel link between mTBI and the increased tendency for neuroticism post-TBI. *Arhgap32* is a CA3 pyramidal cell-specific up-regulated gene in mTBI. It encodes a neuron-associated GTPase-activating protein that may regulate dendritic spine morphology and strength^41^. As it is known that CA3 pyramidal cells play an important role in the relay of information to the rest of the hippocampus, the action of *Arhgap32* may be critical for supporting transmission of information across hippocampal cells. The endothelial-specific gene *Fxyd5* encodes a glycoprotein that functions to enhance chemokine production and inhibit cell adhesion by downregulating E-cadherin^42^. It was upregulated in mTBI. The functions of *Fxyd5* in endothelial cells suggest that it may play a role in neuroinflammation and blood-brain-barrier dysfunction associated with TBI.

### Prioritization of potential targets based on the effects of mTBI on single cells

The aforementioned cell-type specific genes perturbed by mTBI provide unique information about the microcircuits underlying the mTBI pathophysiology. These can be leveraged for the design of therapeutic interventions to target specific cell types driving pathological manifestations if their causal roles in pathogenesis are confirmed. For example, the DG granule cell-specific DEG *Id2* may be targeted for seizure treatment, the CA1 pyramidal cell-specific *Klhl2* can be modulated for personality disorders, the ependymal-specific *Apoe* can be targeted for AD, and the oligodendrocyte-specific *Trf* can be used for iron control. Conversely, identifying DEGs that are affected across multiple cell types by mTBI has the potential to pinpoint the most vulnerable genes that are responsible for the broad symptomology of mTBI. Such genes cannot be retrieved without examining multiple cell types individually to confirm the widespread effect across cell types. Targeting such pan-hippocampal DEGs may offer broader therapeutic effects by normalizing the functions of multiple cell types.

Notably, the gene *Ttr* represents the most robust DEG across hippocampal cell types in that it was a top DEG with increased expression post-mTBI in 7 of the 10 major cell types and 6 of the 8 neuronal clusters, with the highest expression and strongest induction seen in ependymal cells (**Fig. 6b**). *Ttr* encodes transthyretin, the transport protein that carries the thyroid hormone thyroxine, preferentially T4, across the blood brain barrier^43^. It also functions as an amyloid beta scavenger^44^. Moreover, several additional multi-cell type DEGs are related to beta amyloid and AD, including *mt-Rnr2* – encoding the neuroprotective mitochondria factor humanin^45^ which is protective against AD, *Itm2b* – an inhibitor of beta amyloid deposition^46^, and *Apba2* (Mint2) – a stabilizer of amyloid precursor protein APP^47^. These pan-hippocampal DEGs along with some of the cell-type specific DEGs previously discussed strongly implicate pathways related to post-mTBI AD pathogenesis and can be targeted for post-mTBI AD prevention. *Slc17a7* is a pan-neuronal DEG across CA1 pyramidal cells, CA3 pyramidal cells, and DG granule cells. It encodes a sodium-dependent phosphate transporter in neuron-rich regions of the brain and functions in glutamate transport^48^. Interestingly, human genetic polymorphisms in this gene have been associated with recovery time and severity of outcomes after sport-related concussion in humans^49^.

Our single cell data demonstrates *Ttr* as the most robust DEG post-mTBI across hippocampal cell types, which provides a strong rationale for testing *Ttr* as a novel target to modulate mTBI response. As a proof of concept, we examined the possibility that the pan-hippocampal upregulation of *Ttr* might indicate a compensatory need for thyroxine T4, the major brain-specific substrate of transthyretin. Given the strong implication of altered cell metabolism in various cell types discussed above, the critical role of thyroid hormone in regulating metabolism could serve as a platform to mitigate the metabolic crisis after mTBI. We first confirmed, using immunohistochemistry, that Ttr protein was mainly localized to the CA hippocampal subregion (**Fig. 7a**), and the choroid plexus (**Supplementary Fig. 10**) - a reservoir of ependymal cells and important for regulating the passage of substances to the brain through the blood-brain barrier. Acute intraperitoneal injection of T4 post-mTBI protected learning (**Fig. 7b**) and memory (**Fig. 7c**) one week post-mTBI as determined by the Barnes Maze test. This is the first time that T4 treatment was found to mitigate cognitive deficits in a mouse model of concussive injury. A recent study using cortical impact injury suggested a potentially protective action of T4 by measuring a handful of candidate genes related to hypoxia and neurogenesis^50^. In contrast, our study indicates that T4 ameliorates mTBI-induced cognitive dysfunction by engaging the main transporter Ttr, and identified a cascade of genes and pathways associated with cell metabolism through whole transcriptome profiling. We found that T4 treatment affected 121 DEG genes that were also altered by mTBI (**Fig. 7d**). Among the 121 overlapping DEGs, T4 reversed the direction of 93 mTBI signature genes, including *Ttr* (T4 target), *Wdr72* (implicated in cognitive processing speed^51^), and *Tpx2* (protective of neurocyte apoptosis in an AD model^52^) (**Fig. 7e**). To test whether T4 treatment preferentially affects Ttr among the known T4 transporters, we examined the expression pattern of the other T4 transporters in mTBI and in T4 treatment. We found much weaker or lack of alterations in the other transporters by mTBI and/or T4 treatment (**Fig. 7f**). These results suggest that Ttr was indeed the preferential T4 transporter and the effects of T4 treatment was most likely mediated by Ttr. The 93 genes normalized by T4 (**Fig. 7g**) were enriched for hormone response and metabolic pathways (**Fig. 7h**). These results are significant in the face of the metabolic depression caused by TBI, and that T4 reverses the direction of most mTBI signature genes related to metabolism.

**Figure 7.**
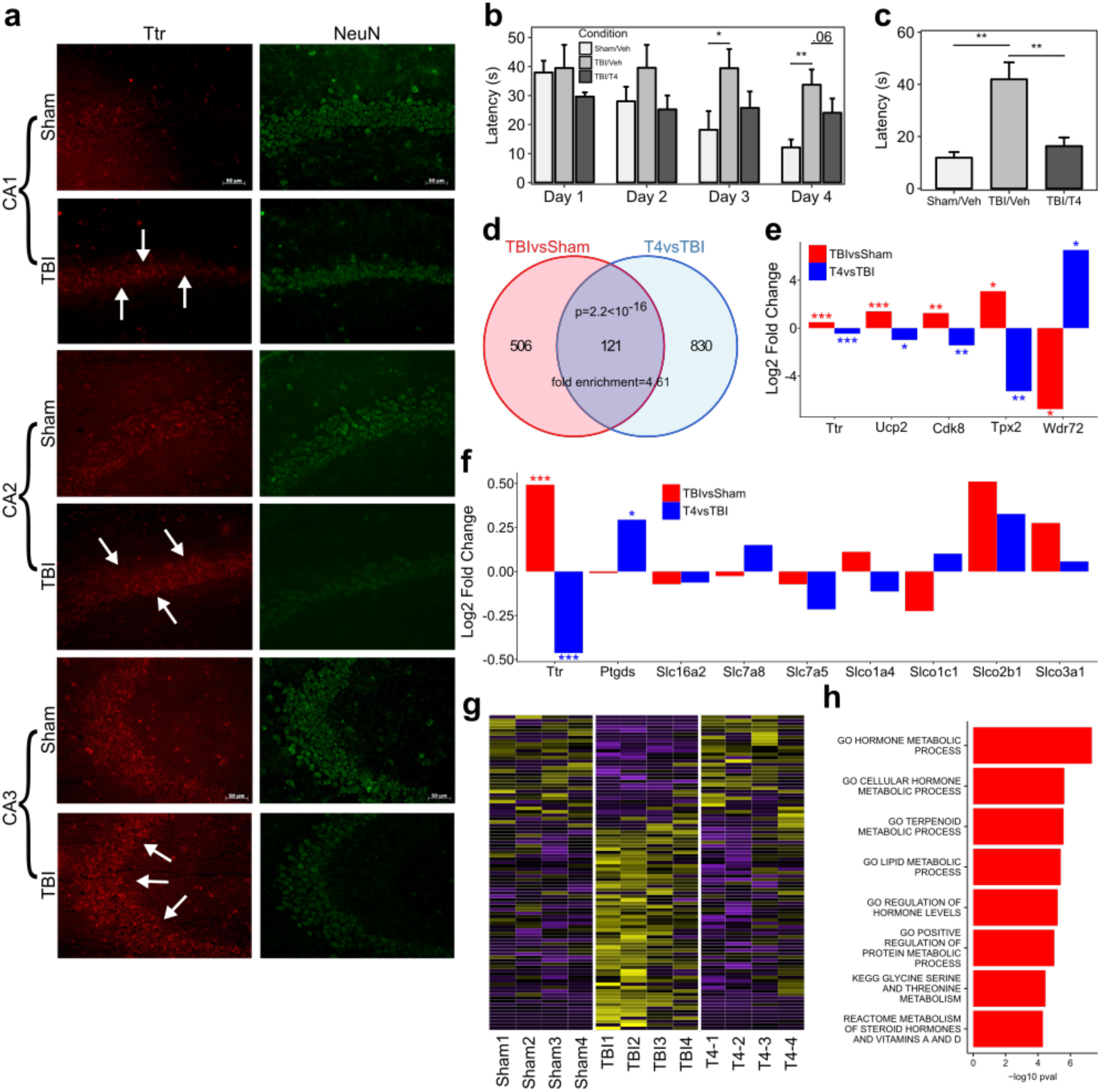
Validation of *Ttr* as a TBI target. (**a**) Immunofluorescence protein expression of Ttr in hippocampal subregions (CA1, CA2, CA3) shows a qualitative increase in staining intensity in Ttr labelled cells in spite of a qualitative reduction in the immunostaining of the neuronal marker NeuN. (**b-c**) T4 treatment corrects TBI-induced learning (**b**) and memory (**c**) deficiency in Barnes Maze test. **(d-e**) Gene expression profiles of T4 treatment experiments. (**d**) TBI and T4 treatment show significant overlap in DEGs. (**e**) Examples of genes reversed by T4. (**f)** Specificity of *Ttr* as the T4 thyroid hormone transporter. (**g**) Heatmap showing T4 reversed the expression patterns of 93 TBI-affected genes. (**h**) Enriched pathways among the 93 DEGs reversed by T4. *p<0.05, **p<0.01, ***p<0.001.

## DISCUSSION

High throughput parallel single cell sequencing analysis proved its utility to unveil key aspects of the molecular and cellular adaptations within the complex cytoarchitecture of the hippocampus in response to mTBI. To our knowledge, this is the first report showing the effects of mTBI on gene regulatory mechanisms in single cells which is crucial to dissect the early events of the mTBI pathology. Our study reveals specific vulnerable cell types including previously undefined cell populations, cell-type specific genes and pathways, and the reorganization of cell-cell interactions, which could be responsible for directing the course of mTBI pathogenesis. Our results on the numerous alterations in cell types, genes, and pathways in response to a single episode of mTBI are remarkable considering that concussive brain injury is difficult to diagnose using conventional neuroimaging examinations^3^. Single cell genomic information is critical for tracing mTBI pathology as the transcriptome can instruct the functions of individual cells and circuits involved in the processing of high order information. Our results suggest the central role of cell metabolism in guiding cell interactions, and we prioritize the cell types and genes involved. The single cell information was indispensable for the prioritization of numerous potential therapeutic targets, including transthyretin, a metabolic regulator affected by mTBI in numerous cell types. The encouraging results from the use of T4 thyroid hormone to intervene *Ttr* and the disrupted metabolic pathways to improve cognitive behavior support the potential of using single cell approaches to identify targets of therapeutic applications. It is important to note that such signals across multiple cell types can only be derived from studying all cell types individually as done in the current study.

The use of functional expression patterns instead of traditional morphological characteristics enabled us to characterize novel cell types, subtypes, and gene markers that are likely more accurate and functionally relevant. The cross-validation of the novel cell markers in independent Allen Brain Atlas datasets supports the reliability of our findings. We acknowledge that we may have missed cell types that are rare or less compatible with the Drop-seq experimental procedures. Nevertheless, the molecular profiles of thousands of single cells of diverse hippocampal cell types under both physiological and pathological conditions represent a rich resource for the neuroscience community to study hippocampus-related processes and diseases.

The transcriptome patterns in individual cell types sensitive to mTBI point to the cellular origins of processes likely guiding mTBI pathogenesis such as metabolic dysfunction, amyloid deposition, and neuronal signaling loss, and contain the gene program regulating and predicting susceptibility to post-mTBI neurological disorders such as AD, PD, PTSD, neuroticism, and epilepsy. The information also has the added potential to guide treatments to improve mTBI outcome by targeting either specific cell types or broad cell interactions. For instance, the fact that genes involved in cell metabolism and amyloid processes were recurring findings across cell types and analytical approaches (cell-type specific DEGs, pan-hippocampal DEGs, pathways, and cell-cell communications) suggests that these are core processes in mTBI pathogenesis. Our single cell study opens new avenues to deconvolute the pathogenic processes involved in mTBI in individual brain cell populations and prioritize cell-type specific (such as *Klhl2* in neurons and *Apoe* in ependymal cells) and pan-hippocampal targets (such as transthyretin and humanin). Concussive brain injury is the most common form of brain injury in sports, domestic, and military settings and has been associated with numerous long-lasting and debilitating neurological consequences. The identified genes, processes, and cell types vulnerable to acute concussive injury will form the foundation for mechanistic studies and for the development of novel therapeutic strategies for mTBI and related neurological disorders.

## Methods

### Animals and mild fluid percussion injury (FPI)

Male C57BL/6J (B6) mice of 10 weeks of age (Jackson Laboratory, Bar Harbor, ME, USA) weighing between 20 and 25g were group housed in cages (n=3-4/group) and maintained in environmentally controlled rooms (22–24 °C) with a 12h light/dark cycle. Mice were randomized to receive either FPI or Sham surgeries. FPI was performed as previously described^53^. Briefly, with the aid of a microscope (Wild, Heerburg, Switzerland), a 1.5-mm diameter craniotomy was made 2.5 mm posterior to the bregma and 2.0 mm lateral (left) of the midline with a high-speed drill (Dremel, Racine, WI, USA). A plastic injury cap was placed over the craniotomy with silicone adhesive and dental cement. When the dental cement hardened, the cap was filled with 0.9% saline solution. Anesthesia was discontinued and the injury cap was attached to the fluid percussion device. At the first sign of hind-limb withdrawal to a paw pinch, a mild fluid percussion pulse (1.5-1.7 atm, wake up time greater than 5 min) was administered. Sham animals underwent an identical preparation with the exception of the lesion. Immediately following response to a paw pinch, anesthesia was restored and the skull was sutured. Neomycin was applied on the suture and the mice were placed in a heated recovery chamber for approximately an hour before being returned to their cages. After 24h, the mice were sacrificed and fresh hippocampal tissue was dissected for use in Drop-seq (n=3/group). All experiments were performed in accordance with the United States National Institutes of Health Guide for the Care and Use of Laboratory Animals.

### Tissue dissociation for Drop-seq

The protocol by Brewer et al.^54^ was used to suspend cells at a final concentration of 100 cells/μl in 0.01% BSA-PBS by digesting freshly dissected hippocampus tissue with papain (Worthington, Lakewood, NJ, USA). Briefly, hippocampi were rapidly dissected from the ipsilateral side of the brain on ice. The hippocampi were transferred into 4 ml HABG (Fisher Scientific, Hampton, NH, USA) and incubated in water bath at 30°C for 8 min. The supernatant was discarded and the remaining tissue was incubated with papain (12 mg in 6 ml HA-Ca) at 30 °C for 30 min. After incubation, the papain solution was removed from the tissue and washed with HABG three times. Using a siliconized 9-in Pasteur pipette with a fire-polished tip, the solution was triturated approximately ten times in 45 sec. Next, the cell suspension was carefully applied to the top of the prepared OptiPrep density gradient (Sigma Aldrich, St. Louis, MO, USA) and floated on top of the gradient. The gradient was then centrifuged at 800g for 15 min at 22 °C. We aspirated the top 6 ml containing cellular debris. To dilute the gradient material, we mixed the desired cell fractions with 5 ml HABG. The cell suspension containing the desired cell fractions was centrifuged for 3 min at 22°C at 200g. We discarded the supernatant, which contained the debris. Finally, the cell pellet was loosened by flicking the tube and the cells were re-suspended in 1 ml 0.01% BSA (in PBS). This final cell suspension solution was passed through a 40-micron strainer (Fisher Scientific, Hampton, NH, USA) to discard debris and cells were then counted.

### Drop-seq single cell barcoding and library preparation

Barcoded single cells, or STAMPs (single-cell transcriptomes attached to microparticles), and cDNA libraries were generated following the drop seq protocol from Macosko et al.^12^ and version 3.1 of the online Drop-seq protocol (http://mccarrolllab.com/download/905). Briefly, single cell suspensions at 100 cells/μl, EvaGreen droplet generation oil (BIO-RAD, Hercules, CA, USA), and ChemGenes barcoded microparticles (ChemGenes, Wilmington, MA, USA) were co-flowed through a FlowJEM aquapel-treated Drop-seq microfluidic device (FlowJEM, Toronto, Canada) at recommended flow speeds (oil: 15,000μl/hr, cells: 4,000μl/hr, and beads 4,000μl/hr) to generate STAMPs. The following modifications were made to the online published protocol to obtain enough cDNA as quantified by a high sensitivity BioAnalyzer (Agilent, Santa Clara, CA, USA) to continue the protocol: 1). The number of beads in a single PCR tube was 4,000. 2). The number of PCR cycles was 4+11 cycles. 3). Multiple PCR tubes were pooled. The libraries were then checked on a BioAnalyzer high sensitivity chip (Agilent, Santa Clara, CA, USA) for library quality, average size, and concentration estimation. The samples were then tagmented using the Nextera DNA Library Preparation kit (Illumina, San Diego, CA, USA) and multiplex indices were added. After another round of PCR, the samples were checked on a BioAnalyzer high sensitivity chip for library quality before sequencing. A cell doublet rate of 5.6% was obtained by running the microfluidic device without the lysis buffer and counting the percentage of cell doublets through three separate runs.

### Illumina high-throughput sequencing of Drop-seq libraries

The Drop-seq library molar concentration was quantified by Qubit Fluorometric Quantitation (ThermoFisher, Canoga Park, CA, USA) and library fragment length was estimated using a Bioanalyzer. Sequencing was performed on an Illumina HiSeq 2500 (Illumina, San Diego, CA, USA) instrument using the Drop-seq custom read 1B primer (IDT, Coralville, IA, USA). Paired end reads were generated using custom read lengths of 24 for read 1 and 76 for read 2 and an 8bp index read for multiplexing. Read 1 consists of the 12bp cell barcode, followed by the 8bp UMI, and the last 4bp on the read are not used. Read 2 contains the single cell transcripts.

### Drop-seq data pre-processing and quality control

Drop-seq tools version 1.12 was obtained from http://mccarrolllab.com/download/922 and the protocol outlined in the Drop-seq alignment cookbook v1.2 (http://mccarrolllab.com/wp-content/uploads/2016/03/Drop-seqAlignmentCookbookv1.2Jan2016.pdf) was followed, using default parameters. Fastq files were converted to BAM format and cell and molecular barcodes were tagged, removing reads corresponding to low quality barcodes. Next, any occurrence of the SMART adapter sequence or polyA tails found in the reads was trimmed. These cleaned reads were converted back to fastq format to be aligned to the mouse reference genome mm10 using STAR-2.5.0c. After the reads were aligned, the reads which overlapped with exons were tagged using a RefFlat annotation file of mm10. A percentage of the Chemgenes barcoded beads which contain the UMIs and cell barcodes were anticipated to have synthesis errors. We used the Drop-seq Tools function DetectBeadSynthesisErrors to quantify the Chemgenes beads batch quality and estimated a bead synthesis error rate of 5-10%, within the acceptable range. Finally, a digital gene expression matrix for each sample was generated where each row is the read count of a gene and each column is a unique single cell. The transcript counts of each cell were normalized by the total number of UMIs for that cell. These values are then multiplied by 10,000 and log transformed. Digital gene expression matrices from the six samples (3 Sham and 3 TBI samples) were combined to create three different pooled digital gene expression matrices for: 1) all Sham samples, 2) all TBI samples and 3) combined Sham and TBI samples. Single cells were identified from background noise by using a threshold of at least 500 genes and 900 transcripts.

### Identification of cell clusters

The Seurat R package (version 1.4.0.1; https://github.com/satijalab/seurat) was used to project all sequenced cells onto two dimensions using t-SNE and density-based spatial clustering (DBSCAN) was used to assign clusters. To further refine the neuronal cell clusters, the BackSPIN software^14^ was used to perform biclustering of the single cells identified to be neuronal cells to further resolve this cell type. Biclustering has been previously demonstrated to differentiate between cell types which cannot be captured by traditional t-SNE-based approaches^14^. BackSPIN was run with default parameters, selecting for the top 2000 most highly variable genes and proceeding with 5 levels of biclustering.

### Identification of marker genes of individual cell clusters

We defined cell cluster specific marker genes from our Drop-seq dataset using a bimodal likelihood ratio test^55^. To determine the marker genes, the single cells were split into two groups for each test: the cell type of interest and all remaining single cells. To be considered in the analysis, the gene had to be expressed in at least 30% of the single cells from one of the groups and there had to be at least a 0.25 log fold change in gene expression between the groups. Multiple testing was corrected using the Benjamini–Hochberg method and genes with an FDR < 0.05 are defined as marker genes. We explored the gene-gene correlation within-group and between-group for each cell type and confirmed the consistency of cell type identification between samples (**Supplementary Fig. 1**).

### Resolving cell identities of the cell clusters

We use two methods to resolve the identities of the cell clusters. First, known cell-type specific markers from previous studies were curated and checked for expression patterns in the cell clusters. A cluster showing high expression levels of a known marker gene specific for a particular cell type was considered to carry the identity of that cell type. Second, we evaluated the overlap between known marker genes of various cell types with the marker genes identified in our cell clusters. Overlap was assessed using a Fisher’s exact test and significance was set to Bonferroni-corrected p<0.05. A cluster was considered to carry the identity of a cell type if the cluster marker genes showed significant overlap with known markers of that cell type. The two methods showed consistency in cell identity determination.

Known markers for major hippocampal cell types and neuronal subtypes were retrieved from Zeisel et al.^14^, Habib et al.^15^, and the Allen Brain Atlas^56^. These markers were sufficient to define all major cell types as well as GABAergic neurons, dentate gyrus (DG), CA1 and CA3 pyramidal neurons.

### Identification of differentially expressed genes between Sham and TBI

Within each identified cell type, Sham and TBI samples are compared for differential gene expression using a bimodal likelihood ratio test^55^. To be considered in the analysis, the gene had to be expressed in at least 30% of the single cells from one of the two groups for that cell type and there had to be at least a 0.25 log fold change in gene expression between the groups. We correct for multiple testing using the Benjamini–Hochberg method and genes with an FDR < 0.05 were used in downstream pathway enrichment analyses (unless explictly noted that a p-value of 0.01 was used instead to retrieve suggestive pathways). Enrichment of pathways from KEGG, Reactome, BIOCARTA, GO Molecular Functions, and GO Biological Processes was assessed with Fisher’s exact test, followed by multiple testing correction with the Benjamini– Hochberg method.

### Comparison of single cell DEGs with those from in silico bulk tissue DEGs

To define the advantages gained by employing single cell sequencing, we simulated bulk tissue gene expression by averaging the gene expression across all TBI single cells and all Sham single cells for each animal. These *in silico* bulk tissue Sham and TBI samples were then compared for differential gene expression using a bimodal likelihood ratio test^55^. FDR<5% was used as the cutoff to determine tissue-level DEGs, which were then compared against those from the single-cell analysis.

### Cell-cell interaction analysis

We assessed cell-cell interactions based on gene-level correlation patterns between any two given cell types (**Fig. 6a**). To infer directionality of the interactions between two cell types, we pointed a cell type whose marker genes encode secreted peptides based on Uniprot information as the source cell type, and then correlated the peptide-encoding marker genes from the source cell type with genes in the target cell type. To deal with the sparse nature of single cell data, we averaged the gene expression of Sham and TBI samples respectively for each cell type. An interaction score is calculated from the sum of the correlation p-values for each peptide, assuming that a peptide from a source cell type with strong correlations with many genes in the target cell type would have a high score and indicate strong interactions. To determine the significance of interaction, we use a permutation approach in which a null distribution is drawn from the interactions scores generated by the correlations between source peptides and target genes where the expression values for each target gene has been shuffled independently. The genes correlated with each peptide were tested for pathway enrichment in KEGG, Reactome, BIOCARTA, GO Molecular Functions, and GO Biological processes to infer the key pathways mediating the interactions.

### Immunohistochemistry

For Klhl2 and Ttr protein immunostaining, Sham and TBI mice were sacrificed and their brains were collected 24 h post-injury. Harvested tissue was mounted with Tissue Tek O.C.T. and then frozen on dry ice and stored at -80 °C. Frozen brains were cryostat sectioned at 10um. Brain sections were fixed in 4% (w/v) paraformaldehyde in phosphate-buffered saline (PBS). Hoechst 33342 (Molecular Probes, Eugene, Oregon, USA) was used for the staining of nuclei. For observation of Klhl2-immunoreactive cells, brain sections were stained using an anti-Klhl2 polyclonal antibody (1:200; Abcam, Cambridge, MA, USA). For the observation of transthyretin (Ttr)-immunoreactive cells, brain sections were stained using an anti-Ttr polyclonal antibody (1:200; Mybiosource, San Diego, CA, USA). For the observation of neuronal nuclei, brain sections were stained using an anti-NeuN polyclonal antibody (1:500; Millipore, Billerica, MA, USA).

### T4 Treatment

L-Thyroxine sodium salt pentahydrate (T4, Sigma Chemical Co., St. Louis, MO, USA) dissolved in saline vehicle (154 nM NaCl) was injected i.p. twice at 1 and 6 h after FPI in the treatment group (n=6 mice) at 1.2 μg/100 g body weight. Control FPI mice (n=6) received vehicle (saline).

### Behavioral tests for T4 treatment experiments

Mice from the Sham, TBI, and T4 treatment groups were tested on the Barnes maze 7 days after injury to assess learning acquisition and memory retention^57^. For learning, animals were trained with two trials per day for four consecutive days, and memory retention was assessed two days after the last learning trial. The maze was manufactured from acrylic plastic to form a disk 1.5 cm thick and 120 cm in diameter, with 40 evenly spaced 5 cm holes at its edges. The disk was brightly illuminated (900 lumens) by four overhead halogen lamps to provide an aversive stimulus to search for a dark escape chamber hidden underneath a hole positioned around the perimeter of a disk. All trials were recorded simultaneously by a video camera installed directly overhead at the center of the maze. A trial was started by placing the animal in the center of the maze covered under a cylindrical start chamber; after a 10s delay, the start chamber was raised. A training session ended after the animal had entered the escape chamber or when a pre-determined time (5 min) had elapsed, whichever came first. All surfaces were routinely cleaned before and after each trial to eliminate possible olfactory cues from preceding animals. After the memory test the animals were sacrificed immediately by decapitation and the fresh hippocampal tissues were dissected out, frozen in liquid nitrogen, and stored in -80°C for bulk tissue RNA-sequencing.

### RNA-seq analysis of T4 treatment experiments

Hippocampal tissues were dissected from Sham, TBI untreated, and TBI T4 treated animals (n=4/group). QuantSeq 3’ libraries were prepared from cDNA samples using the QuantSeq 3’ mRNA-Seq Library Prep Kit (Lexogen, Greenland, NH, USA). Libraries were run on a HiSeq 4000. Reads were aligned with STAR and read counts per gene were generated using the BlueBee platform. Differentially expressed genes between the different groups (Sham, TBI untreated, and TBI T4 treatment) were determined using negative binomial models^58^. DEGs with p < 0.05 were included in gene signatures which were checked for pathway enrichment. DEGs between TBI and Sham were compared for overlap with DEGs between T4 treated mice and untreated TBI mice.

### Accession codes

The NCBI GEO accession number for the Drop-seq data reported in this paper (fastq files and digital gene expression matrices) is GSEXXXXXX.

## Acknowledgements

We thank Dr. Weizhe Hong for advice on neuronal subpopulation analysis and helpful discussions. X.Y. and F.G-P. are funded by R01 DK104363 and R21 NS103088. F.G-P. is funded by R01 NS50465. D.A. is funded by Hyde Fellowship and NIH-NCI National Cancer Institute T32CA201160.

## Contributions

D.A. participated in the experimental design, collected and analyzed sequencing datasets, and wrote the paper. Y.Z., H.R.B., I.S.A., Z.Y., G.Z conducted animal, Drop-seq, and immunohistochemistry experiments, and edited the manuscript. X.Y. and F.G-P. conceived the study, designed and coordinated the study, and wrote the manuscript.

## Supplementary information

### Supplementary Tables

Supplementary Table 1. Markers genes for major hippocampal cell clusters.

Supplementary Table 2. Differentially expressed genes induced by TBI in individual cell clusters.

Supplementary Table 3. Over-represented biological pathways among DEGs between Sham and TBI for individual cell clusters.

### Supplementary Figures

Supplementary Figure 1. Drop-seq library statistics and quality control (QC) features.

Supplementary Figure 2. Cluster specific expression of known cell-type specific markers.

Supplementary Figure 3. CA1 neuron-specific marker gene validation with ISH images from the Allen Brain Atlas.

Supplementary Figure 4. CA3 neuron-specific marker gene validation with ISH images from the Allen Brain Atlas.

Supplementary Figure 5. DG granule cell-specific marker gene validation with ISH images from the Allen Brain Atlas.

Supplementary Figure 6. Multiple neuron cluster-specific marker gene validation with ISH images from the Allen Brain Atlas.

Supplementary Figure 7. CA Subtype 2 neuron cluster-specific marker gene validation and identification of cell type with ISH images from the Allen Brain Atlas.

Supplementary Figure 8. Ependymal-specific marker gene validation with ISH images from the Allen Brain Atlas.

Supplementary Figure 9. Unknown2-specific marker gene validation and identification of cell type with ISH images from the Allen Brain Atlas.

Supplementary Figure 10. Localization of Klhl2 and Ttr protein in the hippocampus.

## References

1. Rohling, M.L., et al. A meta-analysis of neuropsychological outcome after mild traumatic brain injury: re-analyses and reconsiderations of Binder et al. (1997), Frencham et al. (2005), and Pertab et al. (2009). Clin Neuropsychol 25, 608–623 (2011).

2. Girgis, F., Pace, J., Sweet, J. & Miller, J.P. Hippocampal Neurophysiologic Changes after Mild Traumatic Brain Injury and Potential Neuromodulation Treatment Approaches. Front Syst Neurosci 10, 8 (2016).

3. Blennow, K., et al. Traumatic brain injuries. Nat Rev Dis Primers 2, 16084 (2016).

4. Shively, S., Scher, A.I., Perl, D.P. & Diaz-Arrastia, R. Dementia resulting from traumatic brain injury: what is the pathology? Arch Neurol 69, 1245–1251 (2012).

5. Lipponen, A., Paananen, J., Puhakka, N. & Pitkänen, A. Analysis of Post-Traumatic Brain Injury Gene Expression Signature Reveals Tubulins, Nfe2l2, Nfkb, Cd44, and S100a4 as Treatment Targets. Sci Rep 6, 31570 (2016).

6. Meng, Q., et al. Traumatic Brain Injury Induces Genome-Wide Transcriptomic, Methylomic, and Network Perturbations in Brain and Blood Predicting Neurological Disorders. EBioMedicine 16, 184–194 (2017).

7. Redell, J.B., et al. Analysis of functional pathways altered after mild traumatic brain injury. J Neurotrauma 30, 752–764 (2013).

8. Samal, B.B., et al. Acute Response of the Hippocampal Transcriptome Following Mild Traumatic Brain Injury After Controlled Cortical Impact in the Rat. J Mol Neurosci 57, 282–303 (2015).

9. von Gertten, C., Flores Morales, A., Holmin, S., Mathiesen, T. & Nordqvist, A.C. Genomic responses in rat cerebral cortex after traumatic brain injury. BMC Neurosci 6, 69 (2005).

10. Lyeth, B.G. Historical Review of the Fluid-Percussion TBI Model. Front Neurol 7, 217 (2016).

11. Mondello, S., et al. Blood-based diagnostics of traumatic brain injuries. Expert Rev Mol Diagn 11, 65–78 (2011).

12. Macosko, E.Z., et al. Highly Parallel Genome-wide Expression Profiling of Individual Cells Using Nanoliter Droplets. Cell 161, 1202–1214 (2015).

13. van der Maaten, L.J.P. & Hinton, G.E. Visualizing High-Dimensional Data Using t-SNE. Journal of Machine Learning Research 9(Nov), 2579–2605 (2008).

14. Zeisel, A., et al. Brain structure. Cell types in the mouse cortex and hippocampus revealed by single-cell RNA-seq. Science 347, 1138–1142 (2015).

15. Habib, N., et al. Div-Seq: Single-nucleus RNA-Seq reveals dynamics of rare adult newborn neurons. Science 353, 925–928 (2016).

16. Shlosberg, D., Benifla, M., Kaufer, D. & Friedman, A. Blood-brain barrier breakdown as a therapeutic target in traumatic brain injury. Nat Rev Neurol 6, 393–403 (2010).

17. Salehi, A., Zhang, J.H. & Obenaus, A. Response of the cerebral vasculature following traumatic brain injury. J Cereb Blood Flow Metab 37, 2320–2339 (2017).

18. Carlesimo, G.A., et al. Atrophy of presubiculum and subiculum is the earliest hippocampal anatomical marker of Alzheimer’s disease. Alzheimers Dement (Amst) 1, 24–32 (2015).

19. Carlen, M., et al. Forebrain ependymal cells are Notch-dependent and generate neuroblasts and astrocytes after stroke. Nat Neurosci 12, 259–267 (2009).

20. Szabolcsi, V. & Celio, M.R. De novo expression of parvalbumin in ependymal cells in response to brain injury promotes ependymal remodeling and wound repair. Glia 63, 567–594 (2015).

21. Butler, C.R., Boychuk, J.A. & Smith, B.N. Effects of Rapamycin Treatment on Neurogenesis and Synaptic Reorganization in the Dentate Gyrus after Controlled Cortical Impact Injury in Mice. Front Syst Neurosci 9, 163 (2015).

22. Hayes, J.P., Bigler, E.D. & Verfaellie, M. Traumatic Brain Injury as a Disorder of Brain Connectivity. J Int Neuropsychol Soc 22, 120–137 (2016).

23. Bassett, D.S. & Sporns, O. Network neuroscience. Nat Neurosci 20, 353–364 (2017).

24. Fulcher, B.D. & Fornito, A. A transcriptional signature of hub connectivity in the mouse connectome. Proc Natl Acad Sci U S A 113, 1435–1440 (2016).

25. Kanehisa, M. & Goto, S. KEGG: kyoto encyclopedia of genes and genomes. Nucleic Acids Res 28, 27–30 (2000).

26. Croft, D., et al. The Reactome pathway knowledgebase. Nucleic Acids Res 42, D472–477 (2014).

27. Consortium, G.O. Gene Ontology Consortium: going forward. Nucleic Acids Res 43, D1049–1056 (2015).

28. Bye, N., et al. Transient neuroprotection by minocycline following traumatic brain injury is associated with attenuated microglial activation but no changes in cell apoptosis or neutrophil infiltration. Exp Neurol 204, 220–233 (2007).

29. Vespa, P., et al. Metabolic crisis without brain ischemia is common after traumatic brain injury: a combined microdialysis and positron emission tomography study. J Cereb Blood Flow Metab 25, 763–774 (2005).

30. Van Horn, J.D., Bhattrai, A. & Irimia, A. Multimodal Imaging of Neurometabolic Pathology due to Traumatic Brain Injury. Trends Neurosci 40, 39–59 (2017).

31. Sanders, M.J., Sick, T.J., Perez-Pinzon, M.A., Dietrich, W.D. & Green, E.J. Chronic failure in the maintenance of long-term potentiation following fluid percussion injury in the rat. Brain Res 861, 69–76 (2000).

32. Cai, H., et al. Metabolic dysfunction in Alzheimer’s disease and related neurodegenerative disorders. Curr Alzheimer Res 9, 5–17 (2012).

33. Elliott, R.C., Khademi, S., Pleasure, S.J., Parent, J.M. & Lowenstein, D.H. Differential regulation of basic helix-loop-helix mRNAs in the dentate gyrus following status epilepticus. Neuroscience 106, 79–88 (2001).

34. Butovsky, O., et al. Identification of a unique TGF-β-dependent molecular and functional signature in microglia. Nat Neurosci 17, 131–143 (2014).

35. Nathoo, N., Chetty, R., van Dellen, J.R. & Barnett, G.H. Genetic vulnerability following traumatic brain injury: the role of apolipoprotein E. Mol Pathol 56, 132–136 (2003).

36. Matsuda, S., et al. The familial dementia BRI2 gene binds the Alzheimer gene amyloid-beta precursor protein and inhibits amyloid-beta production. J Biol Chem 280, 28912–28916 (2005).

37. Lu, L., Cao, H., Wei, X., Li, Y. & Li, W. Iron Deposition Is Positively Related to Cognitive Impairment in Patients with Chronic Mild Traumatic Brain Injury: Assessment with Susceptibility Weighted Imaging. Biomed Res Int 2015, 470676 (2015).

38. Raz, E., et al. Brain iron quantification in mild traumatic brain injury: a magnetic field correlation study. AJNR Am J Neuroradiol 32, 1851–1856 (2011).

39. Smith, D.J., et al. Genome-wide analysis of over 106 000 individuals identifies 9 neuroticism-associated loci. Mol Psychiatry 21, 749–757 (2016).

40. Soltysik-Espanola, M., et al. Characterization of Mayven, a novel actin-binding protein predominantly expressed in brain. Mol Biol Cell 10, 2361–2375 (1999).

41. Okabe, T., et al. RICS, a novel GTPase-activating protein for Cdc42 and Rac1, is involved in the beta-catenin-N-cadherin and N-methyl-D-aspartate receptor signaling. J Biol Chem 278, 9920–9927 (2003).

42. Ino, Y., Gotoh, M., Sakamoto, M., Tsukagoshi, K. & Hirohashi, S. Dysadherin, a cancer-associated cell membrane glycoprotein, down-regulates E-cadherin and promotes metastasis. Proc Natl Acad Sci U S A 99, 365–370 (2002).

43. Richardson, S.J., Wijayagunaratne, R.C., D’Souza, D.G., Darras, V.M. & Van Herck, S.L. Transport of thyroid hormones via the choroid plexus into the brain: the roles of transthyretin and thyroid hormone transmembrane transporters. Front Neurosci 9, 66 (2015).

44. Oliveira, S.M., Ribeiro, C.A., Cardoso, I. & Saraiva, M.J. Gender-dependent transthyretin modulation of brain amyloid-beta levels: evidence from a mouse model of Alzheimer’s disease. J Alzheimers Dis 27, 429–439 (2011).

45. Bodzioch, M., et al. Evidence for potential functionality of nuclearly-encoded humanin isoforms. Genomics 94, 247–256 (2009).

46. Kim, J., et al. BRI2 (ITM2b) inhibits Abeta deposition in vivo. J Neurosci 28, 6030–6036 (2008).

47. McLoughlin, D.M., et al. Mint2/X11-like colocalizes with the Alzheimer’s disease amyloid precursor protein and is associated with neuritic plaques in Alzheimer’s disease. Eur J Neurosci 11, 1988–1994 (1999).

48. Takamori, S., Rhee, J.S., Rosenmund, C. & Jahn, R. Identification of a vesicular glutamate transporter that defines a glutamatergic phenotype in neurons. Nature 407, 189–194 (2000).

49. Madura, S.A., et al. Genetic variation in SLC17A7 promoter associated with response to sport-related concussions. Brain Inj 30, 908–913 (2016).

50. Li, J., et al. Thyroid hormone treatment activates protective pathways in both in vivo and in vitro models of neuronal injury. Mol Cell Endocrinol 452, 120–130 (2017).

51. Giddaluru, S., et al. Genetics of structural connectivity and information processing in the brain. Brain Struct Funct 221, 4643–4661 (2016).

52. Liang, K., Zhang, J., Yin, C., Zhou, X. & Zhou, S. Protective effects and mechanism of TPX2 on neurocyte apoptosis of rats in Alzheimer’s disease model. Exp Ther Med 13, 576–580 (2017).

53. Wu, A., Molteni, R., Ying, Z. & Gomez-Pinilla, F. A saturated-fat diet aggravates the outcome of traumatic brain injury on hippocampal plasticity and cognitive function by reducing brain-derived neurotrophic factor. Neuroscience 119, 365–375 (2003).

54. Brewer, G.J. & Torricelli, J.R. Isolation and culture of adult neurons and neurospheres. Nat Protoc 2, 1490–1498 (2007).

55. McDavid, A., et al. Modeling bi-modality improves characterization of cell cycle on gene expression in single cells. PLoS Comput Biol 10, e1003696 (2014).

56. Lein, E.S., et al. Genome-wide atlas of gene expression in the adult mouse brain. Nature 445, 168–176 (2007).

57. Barnes, C.A. Memory deficits associated with senescence: a neurophysiological and behavioral study in the rat. J Comp Physiol Psychol 93, 74–104 (1979).

58. Y, D., DW, S., JS, C. & JH, C. The NBP negative binomial models for assessing differential gene expression from RNA-seq. (Stat Appl Genet Mol Biol, 2011).

